# Developmental pyrethroid exposure disrupts molecular pathways for MAP kinase and circadian rhythms in mouse brain

**DOI:** 10.1101/2023.08.28.555113

**Authors:** Jennifer H. Nguyen, Melissa A. Curtis, Ali S. Imami, William G. Ryan, Khaled Alganem, Kari L. Neifer, Nilanjana Saferin, Charlotte N. Nawor, Brian P. Kistler, Gary W. Miller, Rammohan Shukla, Robert E. McCullumsmith, James P. Burkett

## Abstract

Neurodevelopmental disorders (NDDs) are a category of pervasive disorders of the developing nervous system with few or no recognized biomarkers. A significant portion of the risk for NDDs, including attention deficit hyperactivity disorder (ADHD), is contributed by the environment, and exposure to pyrethroid pesticides during pregnancy has been identified as a potential risk factor for NDD in the unborn child. We recently showed that low-dose developmental exposure to the pyrethroid pesticide deltamethrin in mice causes male-biased changes to ADHD- and NDD-relevant behaviors as well as the striatal dopamine system. Here, we used an integrated multiomics approach to determine the broadest possible set of biological changes in the mouse brain caused by developmental pyrethroid exposure (DPE). Using a litter-based, split-sample design, we exposed mouse dams during pregnancy and lactation to deltamethrin (3 mg/kg or vehicle every 3 days) at a concentration well below the EPA-determined benchmark dose used for regulatory guidance. We raised male offspring to adulthood, euthanized them, and pulverized and divided whole brain samples for split-sample transcriptomics, kinomics and multiomics integration. Transcriptome analysis revealed alterations to multiple canonical clock genes, and kinome analysis revealed changes in the activity of multiple kinases involved in synaptic plasticity, including the mitogen-activated protein (MAP) kinase ERK. Multiomics integration revealed a dysregulated protein-protein interaction network containing primary clusters for MAP kinase cascades, regulation of apoptosis, and synaptic function. These results demonstrate that DPE causes a multi-modal biophenotype in the brain relevant to ADHD and identifies new potential mechanisms of action.

**NEW & NOTEWORTHY:** Here, we provide the first evidence that low-dose developmental exposure to the pyrethroid pesticide, deltamethrin, results in molecular disruptions in the adult mouse brain in pathways regulating circadian rhythms and neuronal growth (MAP kinase). This same exposure causes a neurodevelopmental disorder (NDD) relevant behavioral changes in adult mice, making these findings relevant to the prevention of NDDs.

## INTRODUCTION

Neurodevelopmental disorders (NDDs) are lifelong, incurable brain disorders with few biomarkers and few treatments(1). The incidence of NDDs is rapidly rising, with 17% of children in the US now affected, up from 12% in 1998(2). NDDs frequently co-occur as a cluster that includes attention deficit hyperactivity disorder (ADHD), autism, schizophrenia, intellectual disability, communication disorders, and motor and sensory disorders and share a set of common comorbidities, including epilepsy and sleep disorders(3, 4). Recently, evidence of a significant environmental component of NDD risk has been growing. Heritability estimates of autism have been falling in recent years, with large meta-analyses now estimating the environmental component at 38-48%(5). While heritability for ADHD is high, less than 4% of the variance in ADHD diagnosis can be explained by candidate genes, suggesting that the genetic component is being inflated by gene-environment interactions(6–9). The exploration of environmental contributors to NDD risk and their molecular mechanisms has the potential not only for future prevention but also for the discovery of new therapeutics targeting these mechanisms in children and adults.

Pyrethroid pesticides are insect-targeting neurotoxins that are the active ingredients in a wide range of products, including consumer and professional insect sprays and foggers, head lice shampoos, flea collars, and aerosol agents used for mosquito control(10). Pyrethroid pesticides as a class are becoming some of the most widely used in the U.S. due to their low environmental impact and relative safety in adults(10). Pyrethroid use is so widespread that 70-80% of the general US population have detectable pyrethroid metabolites in their blood, indicating an exposure within days(11, 12). Nonetheless, evidence is building that there is a risk for NDD from ambient residential pyrethroid pesticide exposure, particularly during pregnancy. Epidemiology studies have shown that women who lived during pregnancy in areas where pyrethroids were used were at greater risk for their unborn child to later be diagnosed with an NDD(13–15). The level of pyrethroid metabolites in the blood of pregnant women is associated with adverse neural and mental development outcomes in the child(16). Further, pyrethroid metabolite levels in the blood are associated with self-reported ADHD diagnosis(11, 12).

To follow up on these correlational findings in humans, mouse models of pyrethroid exposure have been developed(11, 17–24). These models have established that exposure in mice to the pyrethroid deltamethrin during pregnancy and lactation at doses well below the EPA benchmark dose of 14.5 mg/kg is sufficient to cause broad male-biased changes in the brain and behavior of the resulting offspring. Male mice with developmental pyrethroid exposure (DPE) display hyperactivity, impulsivity, repetitive behaviors, diminished vocalizations, and impaired learning(11, 24). These behavioral changes are in part caused by changes in the striatal dopamine system, including elevated dopamine transporter and dopamine D1 receptor expression, elevated striatal dopamine and dopamine metabolites, and increased peak dopamine release. Other neurological changes have also been observed from developmental and acute exposures, including alterations in levels of voltage-gated sodium channels and brain-derived neurotrophic factor (BDNF), as well as decreased neurogenesis and increased endoplasmic reticulum stress(17–20). These findings suggest that changes in the brain may be much broader than those represented by the largely hypothesis-driven studies performed to date.

A complex biological system such as the brain functions at many biological levels simultaneously, including DNA, mRNA, proteins, and metabolites, each of which can be influenced by a disease or toxin. Any of these changes individually, or multiple multimodal changes across levels, may result in alterations in cellular function that produce a phenotype. RNA sequencing has been used to analyze the pathogenesis of many diseases and shows great potential for the identification of therapeutically significant pathways(25). Further, kinases play a critical role in signal transduction, control most cellular functions, and are abnormally phosphorylated in a wide range of diseases, making kinome array profiling (i.e., functional proteomics) an intriguing avenue for the identification of kinases as potential druggable targets(26–31). Using a combination of high-throughput omics analyses, we can gain a much greater insight into disease mechanisms than with a single omics dataset, allowing for a more in-depth investigation of the underlying biology of the system, potential associations, and other observed effects.

Here, we used split-sample transcriptomics and kinomics to explore, in a hypothesis-free way, the effects of developmental deltamethrin exposure as previously described(24) on the mouse brain at the level of gene transcription and protein activity. Because virtually all prior effects observed in mice were male-biased(11, 24), we chose to focus on male mice. We then combined our transcriptomics and kinomics datasets using multiomics integration analyses to identify multi-modal effects. We hypothesize that indirect developmental deltamethrin exposure via the mother leads to a multi-modal biophenotype in the offspring that may be related to their NDD-relevant behavioral phenotype.

## MATERIALS AND METHODS

### Animals

Experimental subjects were the adult (P56+) male offspring of female C57BL6/J mouse dams. Nulliparous female mice were socially housed in pairs on a 12/12-h light cycle and given water and high-fat breeder food ad libitum. Female pairs were then housed with a male sire until three days prior to birth, at which time they were placed in single housing. The resulting offspring were weaned at 20-22 days of age into same-sex cages and given water and standard mouse diet ad libitum. All procedures were approved by the Emory University IACUC and performed in accordance with the US Animal Welfare Act.

### Study design

The design consisted of two independent groups. Dams were exposed orally to either a low dose of the pyrethroid pesticide deltamethrin (DPE group) or vehicle only (control group) every 3 days throughout pre-conception, pregnancy, and weaning as described below and previously(24). The subjects of the experiment consisted of a single behaviorally naïve adult male offspring selected from each litter.

### Experimental units

Because the exposure occurred in the dam, this study employed a litter-based design, with the dam and the litter together counting as a single experimental unit of N=1. A single behaviorally naïve adult male offspring from each litter was chosen at random for tissue collection. Therefore, the N for each analysis represents both the number of subjects and the number of dams/litters.

### Sample size and elimination criteri

Dams were eliminated from the study if they did not consistently consume the pesticide when offered (as described below) or if they did not give birth to a litter containing at least two male and two female pups within 28 days of the introduction of the sire. This litter composition was needed because at least one male and one female per litter were used separately for behavioral studies(24). A total of 40 dams were offered doses of the pesticide or vehicle (20 per group) as described below. Three dams were eliminated for incomplete consumption; two dams were eliminated for not producing a litter within 28 days; four dams were eliminated for insufficient litter size/composition; and the remaining N=31 dams produced appropriate litters for the study (control N=14, DPE N=17).

### Blinding

Litters were weaned into same-sex cages, which were assigned random numbers that were used for all subsequent purposes. Control and experimental aliquots (described below) were prepared and blindly labeled as A or B. Experimenters remained blind to the identity of the treatments until after data analysis was complete.

### Chemicals

Aliquots of deltamethrin (Sigma, St. Louis, MO) in corn oil (Mazola, Memphis, TN) were prepared at a concentration of 9 mg/mL by dissolving deltamethrin powder in acetone (Sigma), mixing the acetone with corn oil, evaporating the acetone from the mixture overnight in a fume hood, and storing the resulting corn oil in glass vials at −80° C. Vehicle aliquots were prepared using the same volumes of acetone and corn oil. Aliquots were thawed immediately prior to use. Dams were weighed immediately before each feeding and the appropriate amount of corn oil (1 µL/3 g body weight) was mixed with ~100 mg JIF peanut butter (J.M. Smucker, Orville, OH) to achieve a dose of 3 mg/kg.

### Developmental pyrethroid exposure (DPE)

Mice were developmentally exposed to the pyrethroid pesticide deltamethrin as previously described(24). Briefly, dams were given voluntary access to peanut butter containing deltamethrin (3 mg/kg) or vehicle once every 3 days during pre-conception (2 weeks), pregnancy, and lactation. Peanut butter was given during the light cycle, and dams were allowed up to 4 hours for voluntary consumption. To ensure individual dosing, adult cagemates (when present) were separated with dividers, and the peanut butter was put in a high position out of the reach of pups. Dams that failed to consume 90% of the peanut butter during 90% of the feedings were eliminated from the study. Offspring were only exposed to deltamethrin indirectly through the dam. Offspring were weaned from the dam as above and received no additional treatment until adulthood.

### Tissue

Adult male offspring were sacrificed by rapid decapitation and dissected, and the brains were flash-frozen using liquid nitrogen and stored at −80° C until needed. Frozen sagittal half-brain tissue samples (randomized for side) from each of the 31 subjects were pulverized and the powder aliquoted for use in split-sample multiomics.

### Transcriptomics sample processing

RNA extraction was performed on N=31 homogenized half-brain tissue aliquots using the RNeasy Plus Micro Kit (QIAGEN, Germantown, MD). Tissue aliquots were homogenized in 350 µl QIAGEN Buffer RLT Plus using a Qiagen TissueLyser LT and centrifuged for 3 min at maximum speed. Supernatant was removed by pipetting and transferred to a QIAGEN gDNA Eliminator spin column placed in a 2-ml collection tube. The lysate was centrifuged for 30 s at >10,000 rpm to remove DNA contamination from the extract. The flow-through was saved, and 350 µl 70% ethanol was added and mixed well by pipetting. The entire sample was transferred to a QIAGEN RNeasy MinElute spin column placed in a 2-ml collection tube and centrifuged for 15 s at 10,000 rpm. Next, 700 µl Buffer RW1 was added to the MinElute spin column and centrifuged for 15 s at 10,000 rpm. Buffer RPE (500 µl) was added to the MinElute spin column and centrifuged for 15 s at 10,000 rpm. Finally, 500 µl 80% ethanol was added to the MinElute spin column and centrifuged for 2 min at 10,000 rpm to wash the spin column membrane. The spin column was placed into a new 2-ml collection tube and centrifuged for 5 min at full speed to dry the membrane. The spin column was added to a new 1.5-ml collection tube, with 14 µl of RNase-free water added directly to the center of the spin column membrane. RNA was eluted with 1-min centrifugation at full speed. Two aliquots of RNA extract were prepared for each sample: one for RNA concentration and purity analysis and the other to be flash frozen in a dry ice and ethanol mixture. The frozen RNA extracts were then sent overnight on dry ice (via FedEx) to the University of Michigan Advanced Genomics Core (Ann Arbor, MI) to undergo quality control, library preparation and Next Generation Sequencing.

The Total RNA Library was prepared using the Illumina NovaSeq (S4) 300 cycle flow cell and reagent kit (Illumina, San Diego, CA). The high-throughput sequencing strategy included 300 cycles and a read length of paired-end 150 base pairs. An aliquot of extracted RNA for each subject was assayed to determine sample RNA concentration and purity. All samples used in the study passed the RNA quality check via NanoDrop One (Thermo Scientific, Waltham, MA) with a 260/280 ratio of 2, and via Agilent Bioanalyzer with an RNA integrity number (RIN) > 7.

### Transcriptomics quality control

The University of Michigan Advanced Genomics Core provided sample sequences in the form of paired end FASTQ files. Paired end reads were aligned to the mouse reference genome GRCm38 provided by Ensembl (RRID:SCR_002344) using Hisat2 (RRID:SCR_015530)(32). Count data were then generated for the reads aligned to exons and transcripts using the GenomicFeatures (RRID:SCR_016960) and GenomicAlignments (RRID:SCR_024236) packages in R and gene model (GTF file) provided by Ensembl. On average, 31 million unique reads per sample were obtained.

### Genes of interest

We first used the Deseq2 R package (RRID:SCR_015687)(33) to assess the differential expression of all genes in comparisons between the control and DPE groups. Results were reported with both raw and FDR-adjusted p-values. We then used the variancePartition R package (RRID:SCR_019204)(34) to quantify the variation in gene expression attributable to the exposure group. “Genes of interest” were determined by selecting genes that were both (1) differentially expressed between groups (p<0.05 uncorrected) and (2) had >15% variance attributable to the dimension of group.

### Gene set, transcription factor, and kinase enrichment analysis

We performed enrichment analyses using the 65 identified “genes of interest” as input. The web tool Enrichr (RRID:SCR_001575)(35, 36) was used to perform gene set enrichment analysis on the genes of interest using the “Wikipathways 2019 Mouse” pathway database. The web tools ChIP-X Enrichment Analysis 3 (CHEA3, RRID:SCR_023159)(37) and Kinase Enrichment Analysis 3 (KEA3, RRID:SCR_023623)(38) were used to perform transcription factor enrichment analysis and kinase enrichment analysis, respectively. CHEA3 and KEA3 compute enrichment scores for transcription factors and kinases whose protein targets are overrepresented in an input list of genes (or an input list of differentially phosphorylated proteins; see the kinome pipeline below). For both CHEA3 and KEA3, mean rank integrated across libraries was chosen as the level of inquiry.

### Kinomics sample processing

One of the five homogenized sample aliquots was used to measure kinase activity. Lysis buffer was prepared using a 1:100 dilution of M-PER Mammalian Extraction Buffer (Thermo Scientific) to EDTA-free Halt Protease Inhibitor Cocktail (Thermo Scientific) and stored on ice. The sample aliquot was then lysed in the appropriate amount of lysis buffer (3x sample volume) and incubated for 30 min on ice. The lysate was then centrifuged for 15 min at 16,000 g in a pre-cooled centrifuge in a 4°C refrigerator. The supernatant was then extracted and separated into 3 aliquots of equal volume (~15 µl), as freeze thaw cycles can result in loss of kinase activity. One aliquot was set aside for immediate use for protein quantification. The other two aliquots were immediately flash frozen in a dry ice and ethanol mixture and then stored at −80°C until use by the core.

Protein quantification was performed on each sample via a BCA Protein Assay with appropriate standards and reagents. We made standards using BSA stock solution (2 mg/ml), hypertonic buffer, and H_2_O. We added 30 µl of each standard to the designated well in duplicates (n=2) and 2 µl of sample to designated wells with 28 µl of H_2_O in triplicates (n=3). The blank in our assay used BCA buffer +30 µl of H_2_O only. We made a 50:1 mixture of Reagent A to Reagent B, 250 µl which was then added to each well on the 96-well count plate. The plate sat and incubated at room temperature for 30 min. The plate was read at 562 nm with the appropriate plate measurements on SoftMax Pro software. The results were collected and transferred to an Excel document for analysis. Concentrations were determined utilizing the standard curve and level of absorption generated from each sample and used to make the appropriate dilutions prior to transferring samples to the Core Facility.

On the day of processing, the University of Toledo Kinome Array Core Facility performed high-throughput kinase activity profiling on the sample lysates, using the Pamstation12 instrument and the PamChip assay (PamGene, ’s-Hertogenbosch, The Netherlands). The Pamstation12 can be loaded with either a serine/threonine or phosphotyrosine microarray chip, accommodating 3 chips per run. Each chip contains 4 wells, where 4 samples can be loaded. The chips function as microarrays, representing 144 (STK chip) or 196 (PTK chip) reporter peptides that can be phosphorylated by various kinases. Each of the 4 wells in the chip contain well-validated target peptides. Protein kinases is the sample may then phosphorylate target peptides when loaded in the presence of ATP. The Pamstation12 instrument then measures the phosphorylation of these reporter peptides in real time by detecting and imaging fluorescently labeled antibodies at different exposure times and provides the output of each sample’s kinase activity (39). The PamChip arrays were the optimal array for our experiments because they are designed to allow peptide substrates to be deposited at higher concentrations than conventional arrays, thus increasing sensitivity.

The serine threonine kinase (STK) chip can be loaded with a maximum of 2.2 µg of protein per individual array well. The phosphotyrosine kinase (PTK) chip can be loaded with a maximum of 10 µg of protein per individual array well. To meet the appropriate total protein concentrations, the previously frozen extracts were thawed on ice and diluted using ice-cold M-PER buffer concentration (STK total protein = 2.2 µg in 11 µl required volume; PTK total protein = 11 µg in 11 µl required volume). The diluted first-thaw lysate material was then added to 10 µl of MPER buffer and an additional 1 µl of Inhibitor Cocktail, for a total 11 µl of lysate of extract for each sample(39). These extracts were then taken on ice to the University of Toledo Kinome Array Core to be run on the PamGene12 Station. All 31 samples were run on both the STK and PTK chips. Six runs were performed in total, three for STK and the other three for PTK activity profiling.

### Kinomics data analysis

Kinomics data analysis was performed as previously described(29). The BioNavigator software tool was used to preprocess the kinome data, perform image analysis, interpret output measurements, and to visualize, store, and share the results for each of our six runs. The processed data output of BioNavigator was a list of differentially phosphorylated peptide substrates, which was used as the input for subsequent analysis. To maximize our confidence in the prediction of which kinases were upstream of differences in substrate phosphorylation, we used four bioinformatics pipelines to analyze the raw data, each of which relies on its own database and its own mechanism of prediction to identify candidate kinases. The Kinome Random Sampling Analyzer (KRSA) (40) was used as the primary analysis and basis for comparison. The PamGene Upstream Kinase Analysis (UKA), Post-Translational Modification Signature Enrichment Analysis (PTM-SEA)(41), and Kinase Enrichment Analysis Version 3 (KEA3)(38) were used to analyze the same processed data. The Creedenzymatic R package(29) was used to convert the output of each analysis into inclusive percentile rankings, to divide the rankings into quartiles, and to visualize the quartiles in a summary figure. Candidate kinases were identified as “high confidence” if they were detected and appeared in the top quartile of results from all four analyses. Candidate kinases were identified as “medium confidence” if they were detected and appeared in the top quartile of the KRSA results and at least the top two quartiles of the UKA, PTM-SEA, and KEA3 results.

### Kinome Random Sampling Analyzer (KRSA) analysis

KRSA is a validated R package for kinome array analysis and was implemented to identify differential phosphorylation and to predict upstream kinase activity and network modeling(42). This tool allows for a better understanding of kinase interactions and global changes that occur between two states. The output of KRSA is a Z-score value. This value represents the difference between the observed value of a kinase from what you would expect in a random sample. A z-score threshold of 1.5 was used to determine which kinases were considered “hits.” Direction of change in activity was determined for the top 10 kinases by measuring the log_2_ fold change in phosphorylation for each target substrate; assigning each substrate a value change of +1 for a log_2_ fold change (Log2FC) greater than 0.15, 0 for Log2FC between 0.15 and −0.15, and −1 for Log2FC less than −0.15; then averaging the values for all substrates targeted by a kinase. Values above or below zero were used to determine the direction of change. The Creedenzymatic inclusive percentile rankings were calculated based on the absolute value of the Z score.

### Upstream Kinase Analysis (UKA) pipeline

We utilized PamGene’s BioNavigator software to process our datasets through the Protein Tyrosine Kinase Upstream Kinase Analysis (UKA) Knowledge Integration PamApp. The input for the analysis was the raw images from the Pamgene chips. We applied default parameters (scan rank of 4 to 12, 500 permutations) and also incorporated default settings including in vitro/in vivo (1), in silico (PhosphoNET) (1), minimum sequence homology (0.9), minimum PhosphoNET prediction score (300), and a minimum peptide set (3). The calculation of inclusive percentile rankings was based on the absolute value of UKA’s Median Kinase Score.

### Post-Translational Modification Signature Enrichment Analysis (PTM-SEA) pipeline

We employed the publicly accessible repository for the Broad Institute’s Single-sample Gene Set Enrichment Analysis (ssGSEA) and Post-Translational Modification Signature Enrichment Analysis (PTM-SEA)(41) utilizing RStudio Desktop 1.2.5042 (https://rstudio.com) operating within the R 3.3.1 software environment (https://cran.rstudio.com). The input for the PTM-SEA pipeline was the list of differentially phosphorylated peptide sequences. Our data underwent processing through the PTM-SEA pipeline, where we tailored the peptide database to encompass only sequences present on the PamChip, along with peptides having a minimum sequence homology of 0.9. Inclusive percentile ranks were based on the PTM-SEA kinase score.

### Kinase Enrichment Analysis Version 3 (KEA3) pipeline

The KEA3 tool(38) was used as described above for transcriptome analysis, except that the input was a list of differentially phosphorylated substrates. Inclusive percentile rankings were based on the mean rank.

### Network-based multiomics integration of transcriptome and kinome

We used the Kinograte R package(43), which implements an optimized version of the PCSF algorithm(44), to generate an integrated multiomic protein-protein interaction (PPI) network consisting of transcriptomic “hits” (p < 0.05), kinase “hits” (|Z| > 1.5), and interpolated “hidden nodes” connecting them. We assigned node prizes by percentile rank of respective log_2_ fold change (Log2FC) or Z score, and we assigned edge costs by inverse STRING-DB3 interaction confidence(45). Hidden nodes received a node prize of zero by definition. We then ranked PPI network nodes by the average of their node prize and eigencentrality as input for gene-set enrichment analysis using the FGSEA R package (RRID:SCR_020938)(46). We performed gene set over-representation analysis using the PPI network nodes as input to the Enrichr web application(47) with the Gene Ontology database (RRID:SCR_002811)(48). Dysregulated pathways (FDR < 0.05) were used to generate an informed integrated PPI network in order to visualize the 2^nd^ degree interaction network of the top ranked “hub” nodes by eigencentrality, AKT1 and PRKCH.

### Functional interpretation of dysregulated pathways

Dysregulated pathways from FGSEA (FDR < 0.05) were functionally clustered and visualized using PAVER, a meta-clustering method for pathways(49). PAVER finds most representative terms for hierarchically clustered pathway embeddings(50) by selecting whichever term is most cosine similar to its respective cluster’s average embedding. We generated UMAP scatter plots of individual pathways colored and shaped by the cluster they belong to and the experimental comparison they came from, respectively. We then generated heatmaps showing normalized enrichment scores of individual pathways in their identified cluster, arranged by the average enrichment of each cluster.

## RESULTS

### Transcriptome

We performed transcriptomic analysis to compare brain-wide RNA expression in male DPE mice to the expression in vehicle controls. After correction for false discovery rate, differential gene expression analysis identified exactly two genes that differed in their brain-wide expression levels in DPE mice, both of which are canonical clock genes: period circadian regulator 2 (PER2, Log2FC = −0.35, p = 0.014) and circadian-associated repressor of transcription (CIART, Log2FC = −0.42, p = 0.042) (Figure 1A, Supplementary File 1A). Per2 and CIART were under-expressed by 22% and 25%, respectively.

**Figure 1.**
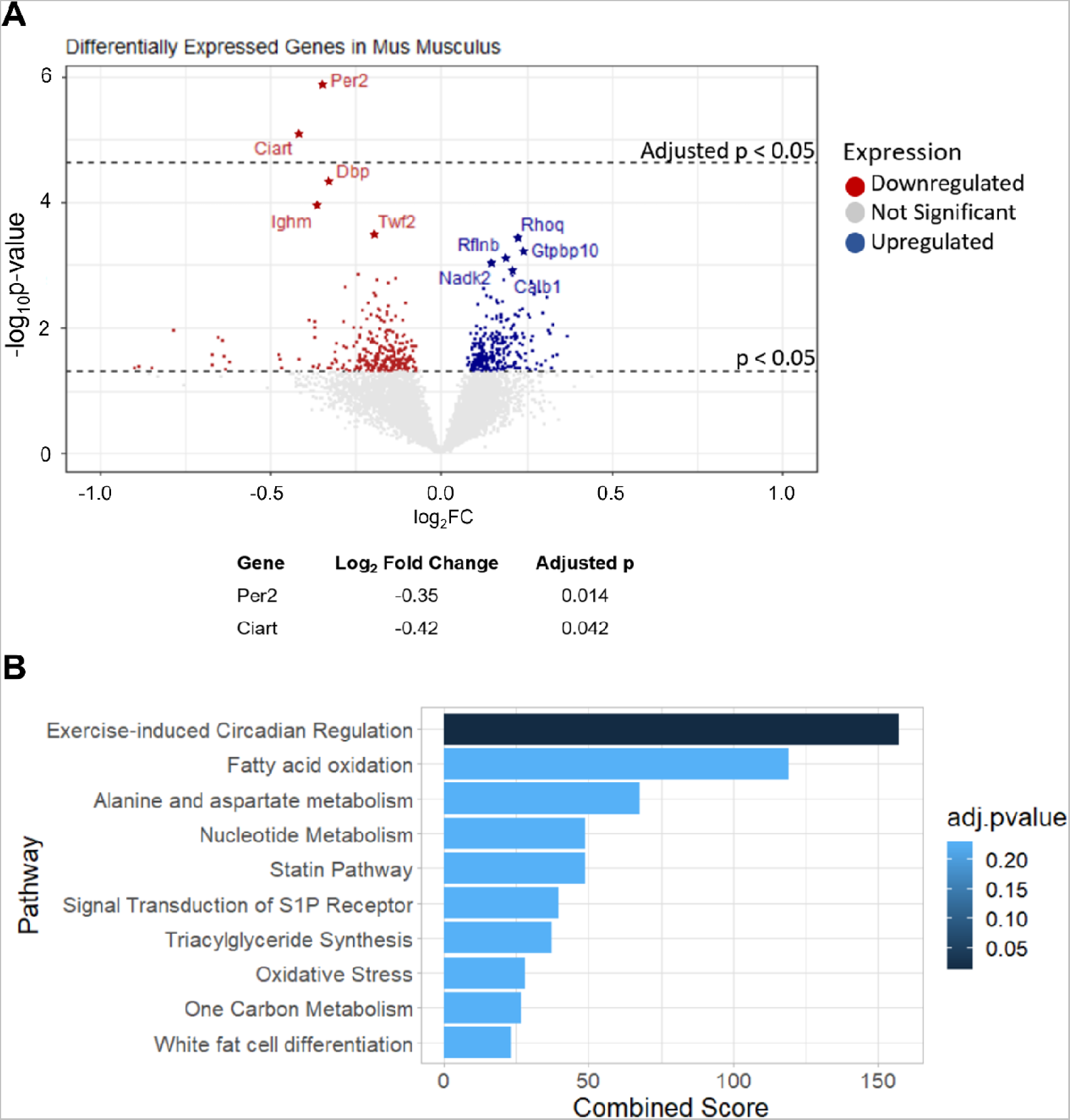
Analysis of transcriptomics data comparing DPE and vehicle-exposed mice. (A) Volcano plot of differential gene expression analysis comparing whole-brain tissue from control and DPE mice, plotted as a function of the log_2_ fold change in gene expression and the inverse log_10_ of the p-value. Dashed lines demarcate raw and adjusted p-values below 0.05. (B) Gene set enrichment analysis on the 65 identified genes of interest. The “Exercise-induced Circadian Regulation” pathway was significantly enriched (adjusted p=0.014).

### Gene sets

Cross-referencing the 607 genes differentially expressed between groups before correction with the 102 genes with >15% of the variance in gene expression explained by exposure produced a list of 65 genes of interest (Supplementary File 1A-C). Gene set enrichment analysis on the 65 genes of interest identified a significant enrichment of genes in the “exercise-induced circadian regulation” pathway (Figure 1B, Supplementary File 1D). Transcription factor enrichment analysis on genes of interest predicted a set of transcription factors upstream of changes in gene transcription, including immediate early genes (FOSB, JUN, NR4A1) and regulators of cell growth (MYC) and neurogenesis (SOX18, PRRX1, TWIST1), among others (Supplementary File 1E). Kinase enrichment analysis predicted kinases upstream of changes in transcription that were related to circadian rhythms (CSNK1A1, CSNK1D, CSNK1E, CSNK2A1), synaptic function (CSNK2A1), cell division/neurogenesis (PRKDC, ATM), and mitogen-activated protein (MAP) kinase/mammalian target of rapamycin (mTOR) cascades (MAPK1/ERK2, EGFR, SRC) (Supplementary File 1F).

### Kinome

We performed kinase activity profiling to detect brain-wide changes in kinase activity in DPE mice. KRSA analysis of differential phosphorylation patterns identified seven kinases with differential activity in DPE mice relative to controls (Z > 1.5), all of which have primary roles in synaptic plasticity (PKCH, PAKB, PKN, MLCK, ERK, BRSK, and PIM), and one of which is the MAP kinase ERK (Figure 2A, Figure S2, Supplemental File 1G). All seven of these kinases had increased activity in DPE mice (Figure 2B, Supplemental File 1H). A summary of four different kinomics analysis algorithms determined that MAPK3/ERK1 and RAF1/c-RAF were high confidence hits on all analyses (Figure 2C, Figure S1).

**Figure 2.**
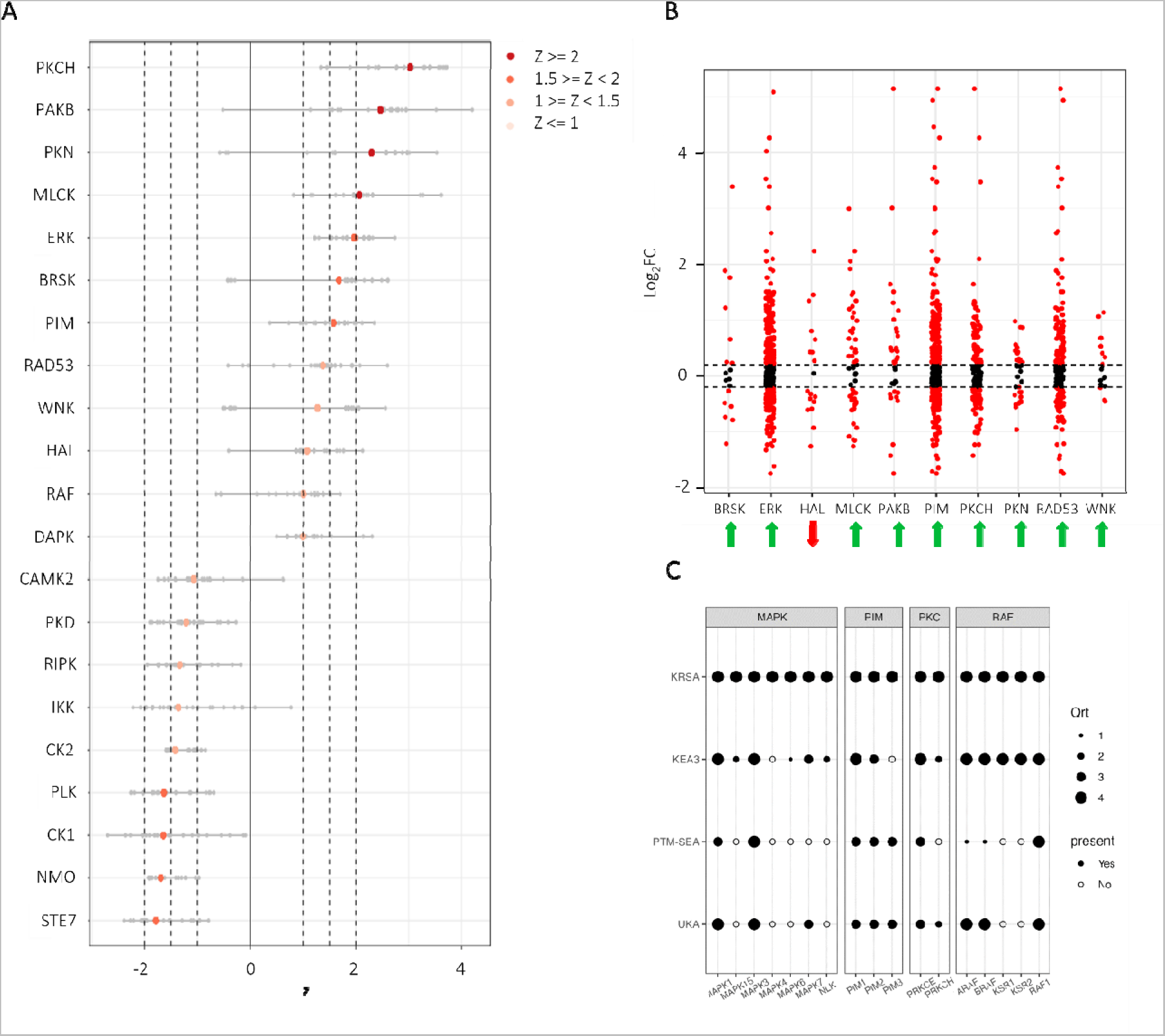
Identification of differentially active serine-threonine kinases identified in whole-brain tissue of DPE mice. (A) Waterfall plot of serine-threonine kinases predicted to be differentially active as a result of differentially phosphorylated peptide substrates on the kinome array chips. (B) Fold-change of phosphorylated substrates related to the top 10 differentially active kinases, showing that most differentially active kinases have increased activity in DPE mice. (C) Summary diagram showing the medium- and high-confidence differentially active kinases as predicted by four different analyses of substrate phosphorylation: KRSA, KEA3, PTM-SEA, and UKA. Dark circles denote kinases whose activity was detected by the analyses. The size of the circles (1-4) denotes the quartile in which the result appears in each analysis, with the 4th quartile being the highest confidence results. Two kinases (MAPK3/ERK1 and RAF1/c-RAF) were high confidence results on all four analyses. Four additional kinases (MAPK1/ERK2, PIM1, PIM2, and PRKCE) were at least medium confidence results on all four analyses.

### Multi-omics Integration

To detect multi-modal changes in molecular pathways, we performed a multi-omics integration of the transcriptome and kinome datasets. Multimodal gene set enrichment analysis returned 521 significantly altered gene sets that clustered into 13 pathways, including MAP kinase cascades, regulation of apoptosis, and synaptic function, among others (Figure 3A-C, Supplementary File 1I). This analysis generated a PPI network with AKT1 and MAPK3/ERK1 as the most central nodes (Figure 3D, Figure S2, Supplementary File 1J, Supplementary File 2).

**Figure 3.**
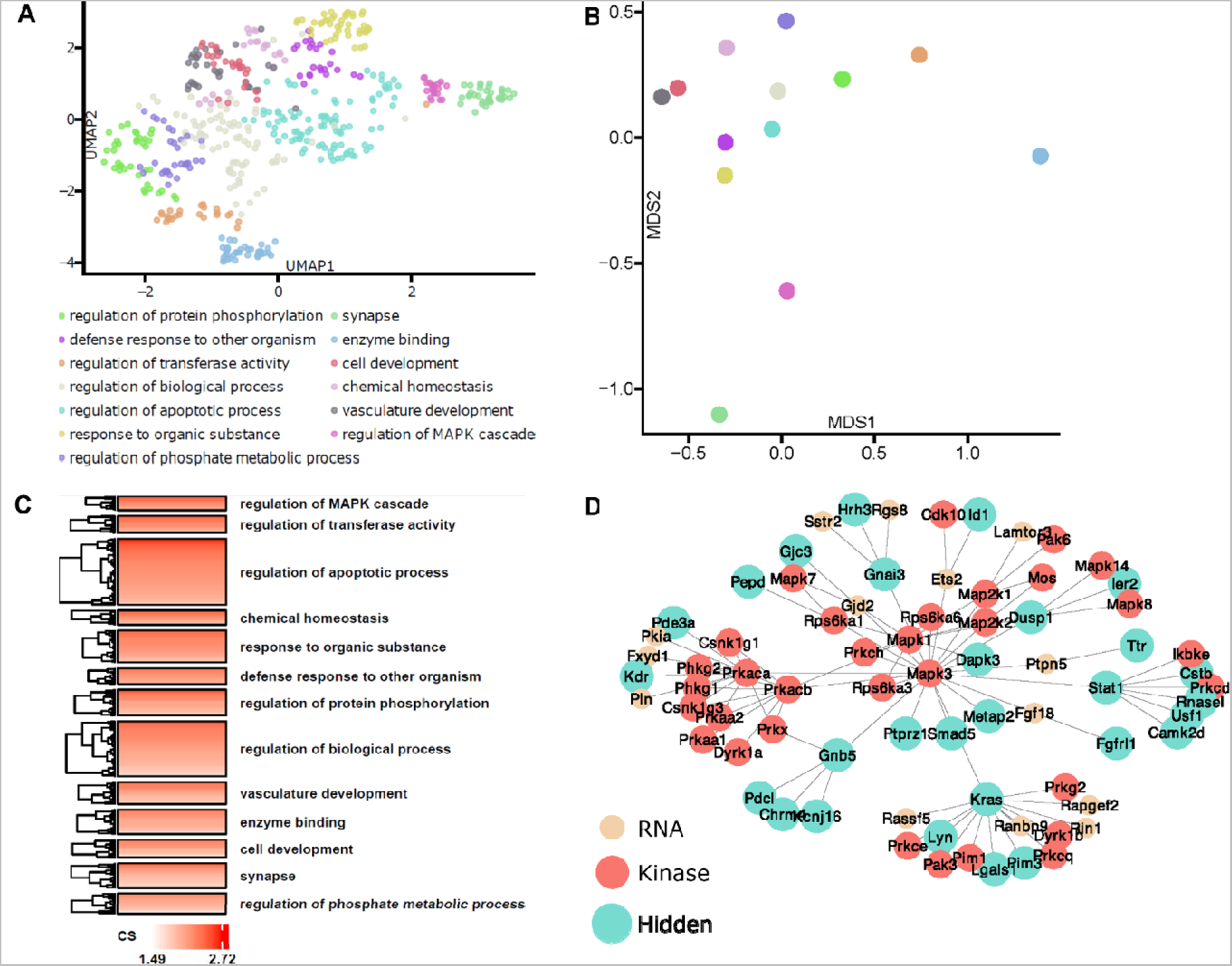
Identification of multi-modal molecular pathways altered in DPE mice using integrative multi-omics. (A) UMAP components analysis of dysregulated molecular pathways identified 13 functionally distinct gene set clusters. (B) MDS components analysis shows the functional relationships between dysregulated gene-set clusters on an interval scale in component space. (C) Pathway cluster heatmap showing the normalized enrichment scores of pathways in the identified gene set clusters, arranged from most (top) to least (bottom) enriched. (D) The 2-degree protein-protein interaction network for the central network hub protein ERK1 (Mapk3).

## DISCUSSION

Developmental exposure to pyrethroid pesticides is a risk factor in humans for autism and ADHD. Using a split-sample multi-omics approach, we detected a multi-modal biophenotype in the brains of mice developmentally exposed to a low level of the pyrethroid pesticide deltamethrin. These multi-modal changes were concentrated in pathways for circadian rhythms, MAP kinase, growth/apoptosis, and synapse function (Figure 4). The brain-wide expression level of genes related to circadian rhythms was altered by developmental exposure. There were also brain-wide increases in the activity levels of a wide range of kinases related to synaptic plasticity, and changes in gene transcription upstream of immediate early genes. Statistical integration of transcriptome and kinome data revealed disruptions in multi-modal molecular pathways clustered around MAP kinase cascades, regulation of apoptosis, and synapses. Overall, we have identified a range of specific targets that may be associated with the neurodevelopmental disorder-related behavioral phenotype observed in these developmentally exposed mice, which includes hyperactivity, repetitive behaviors, decreased vocalizations, and cognitive deficits(24).

**Figure 4.**
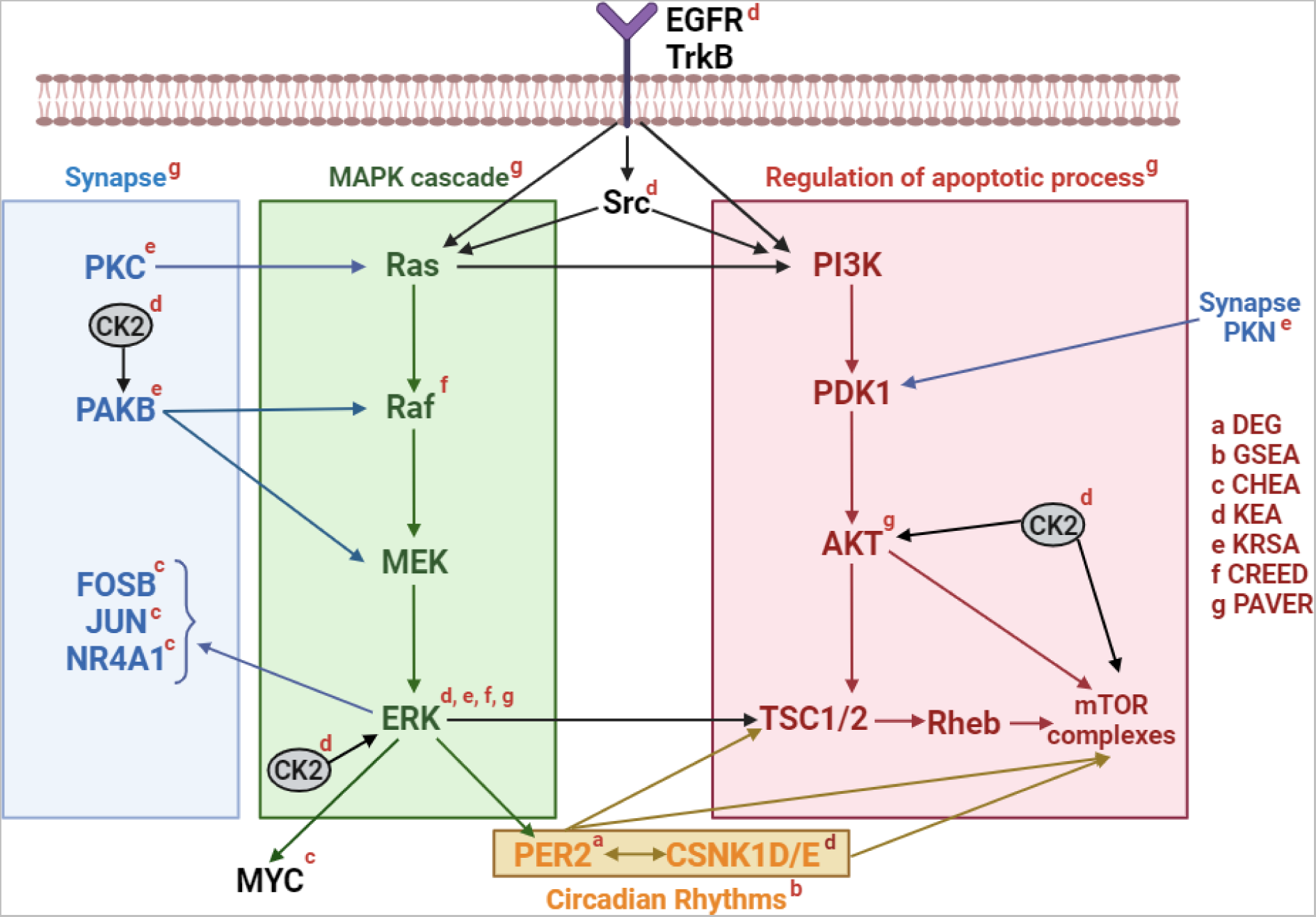
A summary network diagram containing the major findings from multi-omics analysis. DEG: differentially expressed genes. GSEA: gene set enrichment analysis. CHEA: ChIP-X (transcription factor) enrichment analysis. KEA: kinase enrichment analysis. KRSA: kinome random sampling analyzer. CREED: Creedenzymatic analysis. PAVER: Pathway Analysis Visualization with Embedding Representations.

Disruptions in synaptic plasticity and changes in dendritic spines have been implicated in the etiology of autism, ADHD, and neurodevelopmental disorders in general(51–53), and dendritic spine overgrowth is one of the most commonly replicated observations in mouse models of autism risk(54, 55). All seven kinases with increased kinase activity in DPE mice have roles in synaptic plasticity, and synapse function was identified as a significant cluster in the multi-omics network. This broad increase in kinase activity may reflect a biophenotype of dendritic spine overgrowth and/or decreased synaptic pruning, as is seen in autistic patients and some mouse models(3, 53, 54), which needs to be verified experimentally. Protein kinase C (PKC) has a long-established role in long-term potentiation and NMDA regulation(56–58). Protein kinase N, a member of the PKC superfamily, directly interacts with AKT (the central node of our multi-omics network) and may have a specific role in synapse maturation(59, 60). The ERK/MAP kinase cascade regulates transcriptional events related to synaptic plasticity(61); and in addition to being a high-confidence differentially active kinase, ERK substrates were enriched among genes of interest, and ERK was the second most central node in our multi-omics network. PAKB is a PAK family kinase that plays a role in long-term potentiation and dendritic morphology and can regulate MAP kinase(62–64). MLCK, BRSK, and PIM also play roles in various aspects of pre- and post-synaptic signaling and synapse formation(65–67). Finally, transcription factors predicted to be upstream of changes in gene transcription in DPE mice included three immediate early genes (FOSB, JUN, NR4A1) responsive to synaptic transmission. Overall, this collection of observations suggests that alterations in synaptic function and/or dendritic spines are potential biophenotypes in DPE mice that may underlie the learning deficits observed in this model(24).

The MAP kinase and mTOR signaling pathways interact to co-regulate various cellular processes related to cell growth, apoptosis, and adult neurogenesis(68–71) and are linked to neurodevelopmental disorders including autism and epilepsy(64, 72, 73). In addition to their canonical roles in cell growth and differentiation, MAP kinases interact with the mTOR signaling pathway to facilitate dendritic growth and spine formation(68, 74). Both “synapse” and “MAP kinase cascade” were among the multi-modal gene set clusters distinguishing DPE mouse brains from controls, the latter being the most enriched cluster identified. The MAP kinase ERK was identified as a high-confidence hit across four kinome analysis algorithms and was determined to have increased kinase activity in the KRSA analysis. ERK substrates were also disproportionally represented among genes of interest. Two kinases upstream of MAP kinase and mTOR activity, epidermal growth factor receptor (EGFR) and Src, were also predicted to be upstream of transcriptional changes; and ERK is upstream of the immediate early genes FOSB, JUN, and NR4A1. AKT1, a central component of the mTOR pathway, was the most central node of the multi-omics protein-protein interaction network. Overall, these results bolster the hypothesis that MAP kinase and mTOR pathways are disrupted in DPE mice, which may potentially alter synaptic function and/or dendritic spines and could affect pathways regulating apoptosis.

Primary exposure to deltamethrin in adult mice may have a direct effect on adult hippocampal neurogenesis(18). Adult neurogenesis is known to contribute to hippocampal structural and functional plasticity, and is also related to cognition and memory, which are deficient in DPE mice(75, 76). The largest identified multi-modal gene cluster in our data was for the regulation of apoptotic processes, which directly affects adult neurogenesis, in part through the mTOR pathway described above(70, 77–79). While it could be argued that the cluster of gene sets related to apoptosis may be present in our data sets due to the dual role of MAP kinase and mTOR signaling in both apoptosis/neurogenesis and in regulating synapses, MDS component analysis showed that the apoptosis and synapse clusters were quite distant in interval scale component space, suggesting little overlap among the gene sets represented in those clusters. In addition, several of the transcription factors upstream of transcriptional changes in DPE, including PRRX1(80, 81), SOX18(82), and TWIST1(83), are directly involved in regulating Wnt signaling, the classic pathway regulating neurogenesis(84). Therefore, we might expect to discover, on further experimentation, that developmental exposure to deltamethrin in mice via the mother has a similar inhibitory effect on adult neurogenesis in the offspring as acute exposure does in adults.

Disrupted sleep-wake patterns, sleep-onset difficulty, and altered diurnal melatonin patterns have been reported in autism and ADHD(85–90). DPE mice showed transcriptional changes in two canonical clock genes regulating circadian rhythms, PER2 and CIART (also known as CHRONO), which were sufficient to survive genome-wide correction at the whole-brain level. Of note, DBP, the third most differentially expressed gene which did not survive genome-wide corrections, is a transcription factor related to circadian regulation. Transcriptomic genes of interest also significantly clustered in a gene set of circadian regulation, and the circadian rhythm-related transcription factor FOSB was upstream of gene expression changes in DPE. Three circadian rhythm related kinases (casein kinase 1D, casein kinase 1E, casein kinase 1A1) were predicted to be upstream of gene expression changes in DPE, and the differentially active kinase PKC eta also plays a role in circadian rhythms. The suprachiasmatic nucleus traditionally controls the “master clock” for circadian rhythms in the brain, and yet this region is vanishingly small and unlikely to be represented in whole brain homogenate(91). Therefore, the detected changes in clock-related gene transcription and kinase activity are likely to represent pervasive changes in local clocks throughout the brain and may indicate either a primary effect of DPE on local clock function, or a downstream consequence of any number of deficiencies in the function of the suprachiasmatic nucleus. While circadian rhythms were not assessed behaviorally in DPE mice, these mice did show persistent hyperactivity(24). Assessment of circadian rhythms in this model is an important future direction.

A critical strength of this study was the use of an integrative multi-omics approach. Our primary goal was to identify, in the broadest possible sense, the molecular pathways altered by DPE. To accomplish this goal, we used a split-sample approach to maximize the inter-relationship between transcriptome and kinome data. We then employed statistical methods to integrate across data sets in order to examine the emergent multimodal biophenotype across multiple levels of evidence. This holistic, integrative approach to combining multi-omics data highlights the interrelationships of the involved biomolecules and their functions and avoids bias in interpreting parallel data sets(92, 93).

Our gene expression and kinase activity data were generated from samples of homogenized whole brain tissue from mice. Because DPE is a whole-body manipulation, we selected whole-brain tissue as the most appropriate level for initial investigation and did not focus on a specific brain region or cell type. This represents both a significant limitation and a critical strength: any pathways that are disrupted brain-wide in dominant cell types are likely to be good targets for pharmaceutical intervention, while pathways that are not disrupted brain-wide in dominant cell types are unlikely to be represented in our dataset.

The use of a single exposure dose of deltamethrin is a key limitation for this study. Our exposure dose was based on prior studies using this model in which 3 mg/kg developmental exposure was sufficient to cause a broad behavioral and neurological phenotype relevant to neurodevelopmental disorders(11, 18, 24). This dose is well below the lower confidence limit of the benchmark dose (LBMD, 10.1 mg/kg) used for regulatory guidance(10). Nonetheless, a critical next step would be to carefully study the exposure-dependent accumulation of deltamethrin and its metabolites in a variety of tissues and to examine in parallel the dose-dependent effects of exposure on the brain and behavior.

## Perspectives and Significance

Pesticides are widely used around the world, making pesticide exposures during development and adulthood common. Pyrethroid pesticides, while widely considered safe for adults, have been linked to autism and neurodevelopmental disorders in humans and cause related behavioral changes in mice. Here, we show that DPE in mice causes broad changes in molecular pathways in the brain for circadian rhythms, MAP kinase, growth/apoptosis, and synaptic plasticity; and specifically implicate ERK as an important biomarker across multiple levels of analysis. Our findings are suggestive of specific biophenotypes that must be verified experimentally, including dendritic spine overgrowth, altered neurogenesis, and disrupted function of the suprachiasmatic nucleus. These findings point to potential future avenues for medical treatments for neurodevelopmental disorders by targeting these molecular pathways.

## DATA AVAILABILITY

All sequencing data are available on NCBI GEO (GSE241185). All raw kinome array data are available on Zenodo (DOI: 10.5281/zenodo.8283637). All code used to analyze the data is on GitHub repository v1.0 (https://github.com/JamesBurkett7/DPE-disrupts-molecular-pathways-manuscript.git). All metadata, including comprehensive statistical results and protein-protein interaction networks, are available in the supplementary materials.

## SUPPLEMENTAL MATERIAL

Supplemental Figures 1 and 2 can be found in the Supplemental Figures file. Raw data in the form of tables and spreadsheets can be found in Supplemental File 1. The full unedited protein-protein interaction network, in the form of an interactive HTML file, can be found in Supplemental File 2.

## Supporting information

Supplementary File 2 PPI

## ACKNOWLEDGMENTS

We would like to thank S. Daniel of In Scripto, LLC for comments on this manuscript. We acknowledge that Figure 4 was created in Biorender.com. Finally, we acknowledge the ongoing and substantial support provided to our research program by enormous quantities of coffee.

## AUTHOR CONTRIBUTIONS

JPB and GWM designed the exposure study, and JPB and REM designed the multiomics experimental and analytic strategy. All samples were collected in the lab of GWM by JPB and transferred to the lab of JPB. MAC processed all samples, distributed them to the core facilities, and performed the initial analysis of all data. JHN, ASI, KA, and WGR participated in the final analysis of all data and assembly of Figures 1-3 under the guidance of JPB, RS, and REM. NS and JPB created Figure 4. JPB, MAC, and JHN interpreted all results. NS, CNN, and BPK performed background research and assisted in manuscript preparation. KLN provided experimental, administrative, and managerial support. All authors participated in writing the manuscript and approved the final version.

