## Supplementary Figures for "Developmental pyrethroid exposure disrupts molecular pathways for MAP kinase and circadian rhythms in mouse brain"

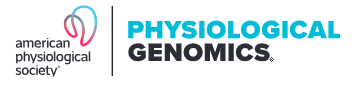

**SUPPLEMENTAL FIGURES FOR:**

Developmental pyrethroid exposure disrupts molecular pathways for MAP kinase and circadian rhythms in mouse brain.

Jennifer H. Nguyen^1†^, Melissa A. Curtis^1†^, Ali S. Imami^1^, William G. Ryan^1^, Khaled Alganem^1,2^, Kari L. Neifer^1^, Nilanjana Saferin^1^, Charlotte N. Nawor^3^, Brian P. Kistler^3^, Gary W. Miller^4,5^, Rammohan Shukla^1,6^, Robert E. McCullumsmith^1,7^, James P. Burkett^1^

^1^ Department of Neurosciences, University of Toledo College of Medicine and Life Sciences, Toledo, OH 43614

^2^ The Medical Cities at the Ministry of Interior, Riyadh, Saudi Arabia (current)

^3^ Department of Medicine, University of Toledo College of Medicine and Life Sciences, Toledo, OH 43614

^4^ Department of Environmental Health, Emory Rollins School of Public Health, Atlanta, GA 30322

^5^ Department of Environmental Health Sciences, Mailman School of Public Health, Columbia University, New York, NY 10032 (current)

^6^ Department of Zoology and Physiology, University of Wyoming, Laramie, WY 82071 (current)

^7^ Neurosciences Institute, Promedica, Toledo, OH 43606

^†^ JHN and MAC should be considered joint first authors.

**This file includes:**

Figures S1-S2

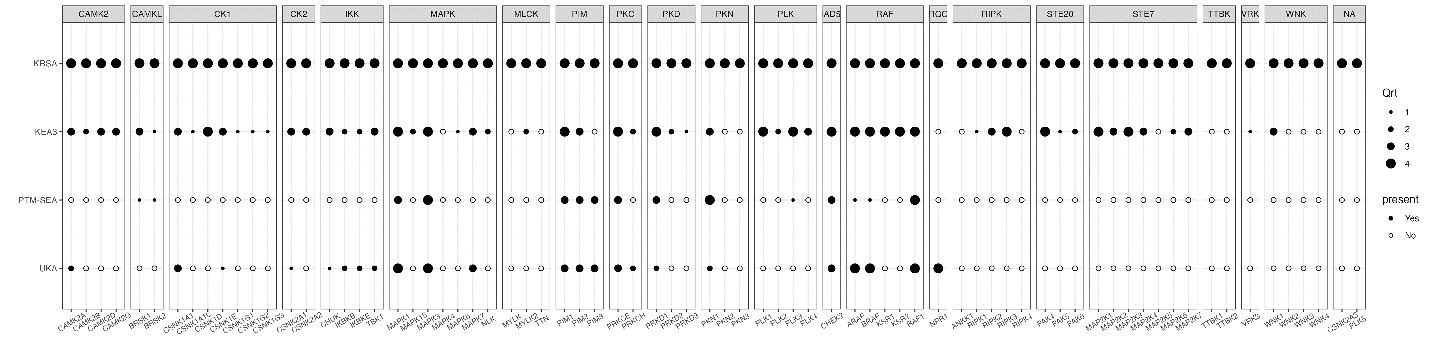

**Figure S1.** The full creedenzymatic result showing differentially active serine-threonine kinases in DPE mouse brains as predicted by four different analyses of kinase chip activity: KRSA, KEA3, PTM-SEA, and UKA. Dark circles denote kinases whose activity was detected by the analysis. The size of the circles (1-4) denotes the quartile in which the result appears, with the 4^th^ quartile being the highest confidence results.

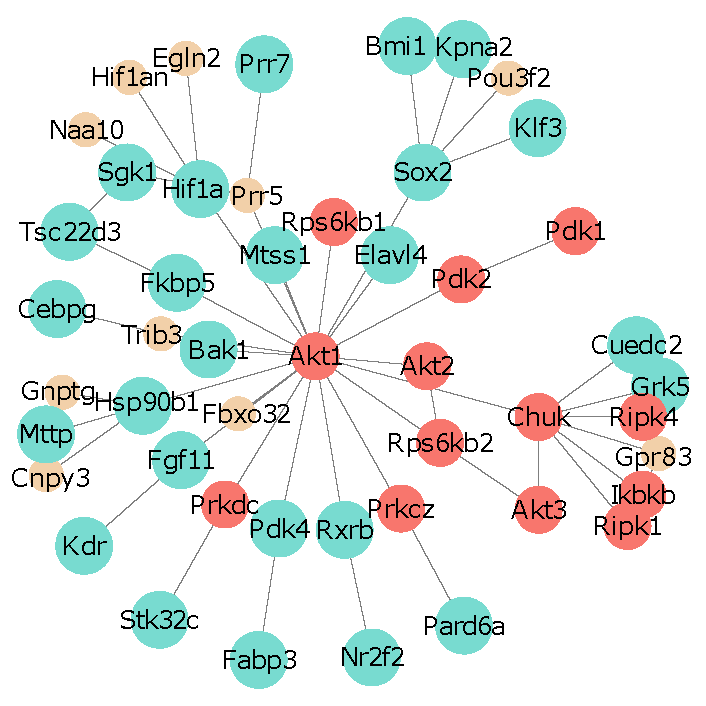

RNA

Kinase

Hidden

**Figure S2.** The 2^nd^-degree protein-protein interaction network for the most central PPI node, AKT1 (Mapk3).
